# Extensive protein dosage compensation in aneuploid human cancers

**DOI:** 10.1101/2021.06.18.449005

**Authors:** Klaske M. Schukken, Jason M. Sheltzer

**Affiliations:** Cold Spring Harbor Laboratory, Cold Spring Harbor, NY 11724

## Abstract

Aneuploidy is a hallmark of human cancers, but the effects of aneuploidy on protein expression remain poorly understood. To uncover how chromosome copy number changes influence the cancer proteome, we have conducted an analysis of hundreds of human cancer cell lines with matched copy number, RNA expression, and protein expression data. We found that a majority of proteins exhibit dosage compensation and fail to change by the degree expected based on chromosome copy number alone. We uncovered a variety of gene groups that were recurrently buffered upon both chromosome gain and loss, including protein complex subunits and cell cycle genes. Several genetic and biophysical factors were predictive of protein buffering, highlighting complex post-translational regulatory mechanisms that maintain appropriate gene product dosage. Finally, we established that chromosomal aneuploidy has an unexpectedly moderate effect on the expression of oncogenes and tumor suppressors, demonstrating that these key cancer drivers can be subject to dosage compensation as well. In total, our comprehensive analysis of aneuploidy and dosage compensation across cancers will help identify the key driver genes encoded on altered chromosomes and will shed light on the overall consequences of aneuploidy during tumor development.

## Introduction

A majority of cancers have chromosome arm gains or losses^1^, a state called aneuploidy. Aneuploidy has been shown to contribute to tumor evolution^2–4^, drug resistance^3,5–9^, and metastatic dissemination^10–12^. High levels of aneuploidy are also associated with poor patient survival^3,13–18^. It is hypothesized that aneuploidy drives tumor development by increasing the dosage of oncogenes (OGs) and decreasing the dosage of tumor suppressor genes (TSGs)^15,19,20^, which may be reflected in the recurrent aneuploid karyotypes found in certain cancer lineages^21^. Despite these findings, aneuploidy itself has also been observed to induce substantial tumor suppressive stresses^10,22,23^. Aneuploid cells have increased metabolic requirements^24–29^, display high levels of senescence^23,30–33^, exhibit significant genomic instability^34,35^, and are sensitive to compounds that interfere with protein folding and turnover^28,36–38^. Aneuploidy-associated stresses may be caused by the deregulation of gene expression, which leads to the imbalanced production of key cellular proteins. How cells adapt to and compensate for these aneuploidy-induced proteome imbalances is a key area of ongoing research.

Genome-wide studies in yeast strains engineered to harbor single extra chromosomes have revealed that copy number gains result in the increased expression of most genes encoded on that chromosome^39,40^. Interestingly, proteomic analysis of these aneuploid yeast strains showed that ∼20% of proteins encoded on extra chromosomes are subject to dosage compensation^39^. This effect is particularly strong for ribosomal subunits and genes that encode subunits of protein complexes^39,41^. However, we have significantly less insight into how aneuploidy shapes the proteome of human cancers. Studies performed on single cancer cell lines harboring a few chromosome gains have suggested that human cells similarly overexpress most proteins on gained chromosomes^24,42–44^, with ribosomes and some protein complex subunits exhibiting dosage compensation^24,41,42,45^. Additionally, while chromosome loss events outnumber chromosome gain events in most tumors^46,47^, the cellular effects of chromosome losses in human cancers have not been comprehensively explored^48^.

Currently, we lack a genome-wide understanding of the effects of aneuploidy on protein expression in cancer. Additionally, the factors that determine which genes are subject to dosage compensation are unknown. Here, we examine a cohort of 367 human cancer cell lines with known chromosome copy numbers, RNA expression levels, and protein expression levels^49^. We demonstrate pervasive patterns of dosage compensation across aneuploid cells, including in particular protein complex subunits, spliceosome components, and cell cycle genes. Sensitivity analysis identifies a set of features shared across buffered proteins, including high levels of ubiquitination and protein-protein interactions. Finally, we find that many key oncogenes and tumor suppressors are resistant to aneuploidy-mediated dosage changes, complicating the hypothesis that aneuploidy-induced alterations in these genes drive tumorigenesis.

## Results

### Mean protein expression increases upon chromosome gain and decreases upon chromosome loss

We examined data from a collection of 367 human cancer cell lines from the Cancer Cell Line Encyclopedia (CCLE) with matched DNA copy number, RNA expression and protein expression data^49–54^ (Supplementary Data 1). We used a recently-described dataset in which each chromosome arm in a cell line was classified as “lost”, “gained”, or “neutral”, relative to that cancer cell line’s basal ploidy^55^. We calculated mean RNA and protein expression differences along each chromosome arm between cell lines with neutral ploidy for that arm and cell lines in which that arm was either gained or lost (Supplementary Data 2). We found that average RNA and protein expression increases upon chromosome arm gain and decreases upon chromosome arm loss (Figure 1A-B). For instance, in cancer cell lines with chromosome 5q gains, the genes located on chromosome 5q tend to exhibit increased expression, while in cell lines with 5q losses, the genes located upon 5q tend to exhibit decreased expression relative to cell lines in which 5q exhibits a neutral ploidy (Figure 1C). While mean gene expression differences correlated with chromosome copy number, we observed a large range of expression differences within the genes located on single aneuploid chromosomes (Figure 1C).

**Figure 1.**
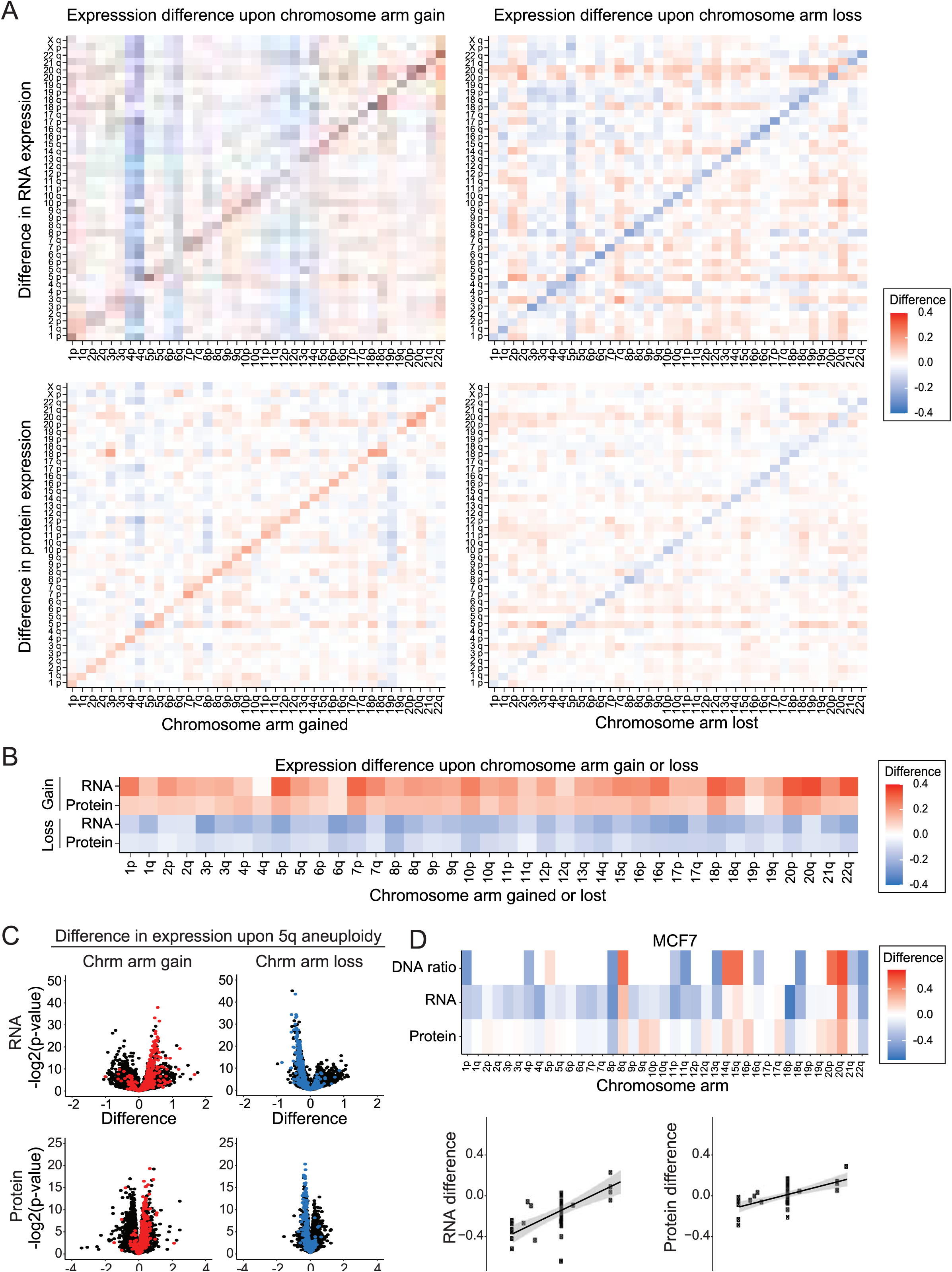
Change in RNA and protein expression upon chromosome arm gain or loss. A) Heatmaps displaying the difference in RNA expression (top) or protein expression (bottom) between cell lines with a chromosome arm gain (left) or chromosome arm loss (right) compared to cell lines with neutral ploidy for that arm. The mean RNA or protein expression differences per chromosome arm are displayed. B) Heatmaps displaying the same analysis as in A, but only for each aneuploid chromosome. C) Volcano plots displaying the difference in RNA expression (top) and protein expression (bottom) vs. the p-value, per gene, between cells with chromosome 5q gain (left) and cell with chromosome 5q loss (right) compared to cell lines with neutral ploidy for chromosome 5q. Genes encoded on chromosome 5q are indicated in red (upon gain) or blue (upon loss), while all other genes are indicated in black. D) Heatmap of the DNA ratio, RNA expression difference, and protein expression difference, per chromosome arm, in MCF7 cells relative to cells neutral for each chromosome arm (top). Scatterplots showing the relationship between the DNA ratio and the RNA or protein expression differences (bottom). Linear regression with 95% confidence intervals is plotted against the data.

The effects of aneuploidy on gene and protein expression were also apparent within individual cancer cell lines (Figure 1D, S1A-E). For instance, in highly-aneuploid cancer cell lines like MCF7 (Figure 1D) and NCIH838 (Figure S1E), genes that were encoded on lost chromosomes tended to be expressed at lower levels, and genes that were encoded on gained chromosomes tended to be expressed at higher levels, relative to the mean expression of those gene in cell lines neutral for that chromosome arm.

Notably, we found that aneuploidy-driven effects on gene expression were more pronounced at the RNA level than at the protein level. For chromosome gains, we found that the average transcript encoded on that chromosome increased by an average of 21%, while protein expression increased by only 12%. A similar pattern was apparent upon chromosome loss, which we found to decrease RNA expression by 24% and protein expression by 9%. These differences suggest that both transcriptional and post-transcriptional dosage compensation can buffer the effects of aneuploidy on gene expression in cancer.

### Ploidy gains buffer the effects of aneuploidy

We next investigated whether the difference in gene expression associated with aneuploidy was affected by a cell line’s basal ploidy. Our dataset of human cancer cell lines consisted of 50% (185/367) near-diploid cell lines and 37% (137/367) near-triploid cell lines, with the remaining 13% spread across other ploidies. We therefore analyzed the near-diploid and near-triploid cell lines separately, and we excluded the other ploidies due to the low number of cell lines available. We found a significant effect of aneuploidy on RNA and protein expression in both near-diploid and near-triploid backgrounds. In both ploidies, the relative expression differences were reduced compared to the relative DNA copy number change (Figure S1F). There was a moderately smaller effect of aneuploidy on gene expression in near-triploid cell lines compared to near-diploid cell lines. For example, in near-diploid cell lines, chromosome gains (corresponding to a 50% increase in DNA copy number) resulted in a 14% increase in protein expression, while in near-triploid cell lines, chromosome gains (corresponding to a 33% increase in DNA copy number) resulted in a 10% increase in protein expression. These results demonstrate that aneuploidy shapes the gene expression landscape in both diploid cancer cell lines and cancer cell lines with higher ploidies, and that ploidy gain has a small but noticeable buffering effect on aneuploidy-driven gene expression changes.

### mRNA changes strongly influence but do not fully explain the effects of aneuploidy on protein expression

Next, we sought to understand the effects of aneuploidy on the expression of individual genes. We compared expression levels between cell lines in which a gene of interest was present on a lost, neutral, or gained chromosome, and we observed several different aneuploidy-driven expression patterns. The expression levels of some genes were significantly increased upon gain of the corresponding chromosome arm and significantly decreased upon loss of that chromosome arm, at both RNA and protein levels (e.g., SMCHD1, Figure 2A, 9% of genes). Other genes significantly changed at the RNA level, but not at the protein level (e.g. NDUFV2, Figure 2B, 13% of genes). Many genes had no significant change in expression upon chromosome arm gain or loss (e.g. CDKN1A, Figure 2C, 19% of genes). Finally, the remaining 59% of genes exhibited more complex and/or variable expression patterns. For instance, GOLGA2 protein expression significantly increased upon chromosome gain, but expression levels of this protein were not significantly affected by chromosome loss (Figure 2D).

**Figure 2.**
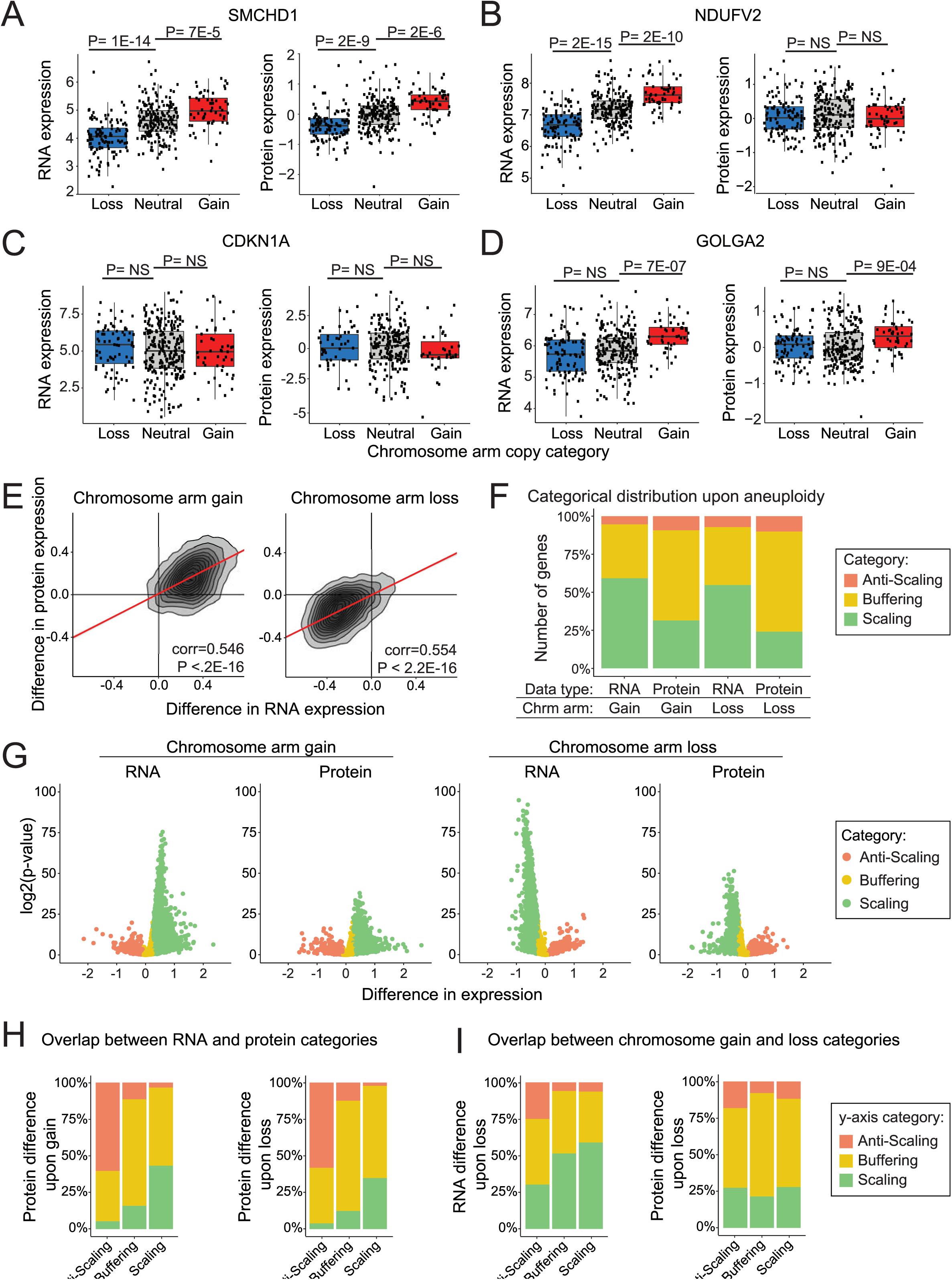
Protein expression differences are frequently buffered upon chromosome gain or loss. A-D) Normalized RNA and protein expression levels of the indicated genes (SMCHD1, NDUFV2, CDKN1A, and GOLGA2) are displayed for cell lines in which the chromosome that that gene is encoded on is either lost, neutral, or gained. Boxplots display the 25th, 50th, and 75th percentile of the data, while the whiskers indicate a 1.5 interquartile range. P-values were calculated using two-sided t-tests. E) A density plot comparing the difference in RNA expression vs. the difference in protein expression levels upon the gain (left) or loss (right) of the chromosome arm the gene is located on. Pearson correlation coefficient and p-value are displayed. Linear regressions (red) with 95% confidence intervals are displayed. F) The percent of RNAs and proteins that fall into each difference category upon chromosome gain and loss are displayed. G) Volcano plots displaying the difference in RNA expression or protein expression upon chromosome arm gain (left) or chromosome arm loss (right) vs. the p-value for each gene. Genes are color coded based on a categorical distribution as either scaling, buffering, or anti-scaling. Categorical cutoff points are at -0.1 and 0.25 for chromosome gain, and at -0.25 and 0.1 for chromosome loss. H) Bar graphs displaying the percent of genes in each RNA difference category (x axis) whose corresponding proteins fall into each of the indicated expression categories upon chromosome gain (left) and loss (right). I) Bar graphs displaying the percent of RNAs in each RNA difference category upon chromosome gain (x axis) that fall into each of the indicated expression categories upon chromosome loss (y axis) (left). Percent of proteins, per protein difference category upon chromosome gain (x axis), that fall into each expression category upon chromosome loss (y axis) (right).

We investigated the degree to which these protein expression changes were determined by the effects of aneuploidy on mRNA expression. We found that RNA expression differences exhibit a significant but imperfect genome-wide correlation with protein expression differences upon chromosome arm gain (Pearson’s correlation coefficient = 0.546, P < 2.2E-16) and upon chromosome arm loss (Pearson’s correlation coefficient = 0.554, P < 2.2E-16) (Figure 2E). These correlations indicate that, while protein expression differences are strongly driven by RNA expression differences, additional factors influence the expression of proteins encoded on aneuploid chromosomes.

### Certain genes exhibit consistent dosage-compensation upon chromosome gain or chromosome loss

We categorized genes as either “scaling”, “buffered” or “anti-scaling”, based on their expression changes upon chromosome gain or loss (Figure 2F-G). Genes that increased (>0.25 log2-fold) with chromosome gain, or decreased (< -0.25 log2-fold) with chromosome loss, were classified as “scaling” with aneuploidy. Genes that were less affected by aneuploidy than expected (−0.1 to 0.25 log2-fold difference upon gain and -0.25 to 0.1 upon loss) were classified as “buffered”. Finally, genes whose expression changed in the opposite direction of what was expected upon chromosome gain or loss (< -0.1 log2-fold difference upon gain or >0.1 log2-fold difference upon loss) were classified as “anti-scaling”. At the RNA level, a majority of genes scaled with aneuploidy, while at the protein level, a majority of proteins were classified as buffered (Figure 2F). For instance, upon chromosome gains, 59% of genes scaled at the RNA level, while 59% of genes exhibited buffering at the protein level. In contrast, only ∼5% of RNA and ∼10% of proteins exhibited an anti-scaling expression pattern.

We observed a significant correlation between RNA and protein difference categories upon chromosome gain (chi-squared = 2447, P < 2E-16) and loss (chi-squared = 2580, P < 2E-16, Figure 2H). For example, upon chromosome gain, 73% of genes that were buffered at RNA level were also buffered at the protein level. Next, we investigated whether genes that were buffered upon loss were more likely to be buffered upon gain. We found that there was a small but significant correlation between gene expression categories upon chromosome gain and loss at the RNA level (chi-squared = 359, P < 2E-16) and at the protein level (chi-squared = 182, P < 2E-16, Figure 2I). For example, 63% of proteins buffered upon chromosome loss were also buffered upon chromosome gain. These results suggest that certain genes exhibit consistent expression patterns, independent of whether the chromosome that they are encoded on is gained or lost.

### Protein complex subunits and cell cycle genes tend to be buffered upon both chromosome gain and chromosome loss

We investigated whether genes with shared functions displayed similar expression patterns when encoded on aneuploid chromosomes. Gene Ontology Enrichment Analysis (GOEA) was used to identify biological terms enriched among scaling, buffered, and anti-scaling genes (Figure 3A, S2A-B, Supplementary File 1-2)^56^. To visualize these expression patterns, we generated density plots of the expression differences of subsets of genes that were encoded on aneuploid chromosomes (Figure 3B). Consistent with previous results^24,26,41,42^, we observed that ribosomal genes, RNA processing genes, spliceosome components, and other genes that encode protein complex subunits were enriched among genes that were buffered upon chromosome gain. Interestingly, we also found that these proteins tended to be buffered upon chromosome loss, and decreased in expression less than other proteins encoded on lost chromosomes. We observed that, on average, protein complex subunits (CORUM genes) increase 21% less than other genes upon chromosome gain and decrease 13% less than other genes upon chromosome loss. Additionally, we found that a majority of genes encoding cell cycle components exhibited protein buffering upon chromosome gain and loss (64% and 71% of proteins, respectively). These findings suggest that dosage compensation is not simply a response to protein overproduction from extra chromosomes, and cancers have the ability to rebalance proteins and protein complex subunits when those genes are encoded on lost chromosomes as well.

**Figure 3.**
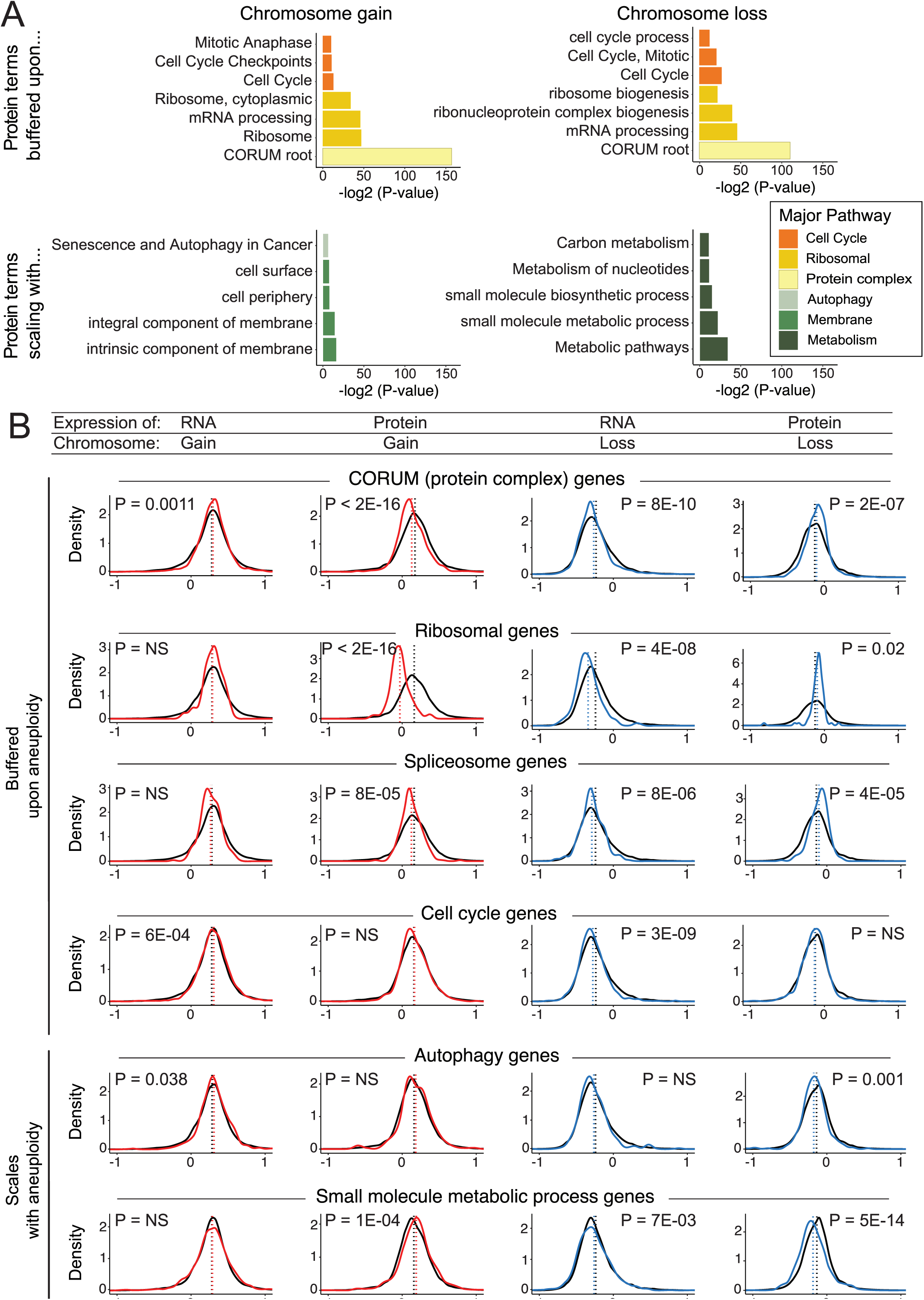
Specific gene groups tend to exhibit buffering or scaling upon chromosome gain and loss. A) Bar graphs displaying the biological terms enriched in proteins buffered or scaling upon chromosome gain or loss, categorized by the major overarching pathway. The complete GO term lists are included in Supplementary Files 1-2. B) Density graphs displaying the difference in RNA or protein expression per gene group upon chromosome gain (red) or loss (blue). The difference in expression for all other genes is shown in black. The mean difference per gene group is indicated by the dotted lines. P-values represent two-sided t-tests between the indicated gene groups and the background set of all other genes.

While many biological pathways were enriched among buffered proteins we found that few GO terms were enriched among buffered RNAs (Figure S2B, Supplementary File 1-2). Indeed, mRNAs coding for protein complex subunits and cell cycle genes were found to be moderately enriched among RNAs that scaled with chromosome copy number, rather than among the buffered subset. This lack of dosage compensation at the RNA level indicates that translational or post-translational regulation, rather than transcriptional regulation, drives the pervasive dosage compensation of buffered proteins and protein complex subunits that we have observed in aneuploid cancer cells.

While a significant percentage of proteins scale with chromosome gain (31%) and loss (24%), we found that few GO terms were enriched among the “scaling” protein category (Figure 3A). The notable exception was metabolic pathways, especially small molecule metabolic pathways, which were found to scale in expression with chromosome loss (Figure 3A-B). There was also a slight enrichment of membrane-associated genes and autophagy genes among genes that scale with chromosome gain (Figure 3A-B).

Only 5-10% of genes exhibited an expression difference in the opposite direction of the DNA copy number change, which we classified as “anti-scaling”. We found that genes associated with the extracellular environment and cell adhesion were enriched among anti-scaling genes at both the RNA level and the protein level and upon chromosome gain and chromosome loss (Figure S2B-C). This may reflect the wide variability in the expression within these gene sets (discussed in more detail below).

In total, we observed that specific gene groups and pathways exhibit similar expression patterns when present on aneuploid chromosomes, including both gained and lost chromosomes. These shared regulatory patterns may arise as a result of genetic and biophysical features that are shared among these genes (discussed in more detail below).

### Patterns of dosage compensation are conserved between aneuploid cancers, trisomic primary cell lines, and disomic yeast

We hypothesized that the expression differences we observed between cell lines that have or lack certain aneuploidies represent a consequence of that chromosome copy number change. Alternatively, these expression patterns could be dictated by a cell line’s lineage or genetic background. If these patterns of scaling and buffering represent a fundamental consequence of aneuploidy, then we would expect to observe similar expression patterns in other aneuploid cell lines. To investigate this possibility, we examined published transcriptome and proteome data from stably-aneuploid human cell lines^24^, Down syndrome fibroblasts^57,58^, and aneuploid yeast^39^.

Previously, Stingele et al.^24^ created aneuploid cell lines by introducing one or more extra chromosomes into chromosomally-stable colon cancer cell lines and non-transformed epithelial cell lines. They then determined the effects of these chromosome gains on gene expression relative to their near-euploid parental cell lines. We examined the difference in protein expression of proteins located on chromosome 5 across four cell lines engineered to gain chromosome 5. We found that there was a highly-significant correlation between the expression difference of proteins encoded on chromosome 5 among CCLE lines with chromosome 5 gains and the expression of those same proteins in the engineered cell lines (Figure S3A) (corr = 0.431, P = 4E-16). Next, we examined protein differences in fibroblasts from Down syndrome patients (with Trisomy 21) normalized to matched euploid fibroblasts. We observed a strong and significant correlation between protein expression differences of proteins located on chromosome 21 in Down syndrome patients and in CCLE cell lines with gains of chromosome 21 (Figure S3B) (corr = 0.539, P = 3E-04).

To investigate whether these patterns of protein dosage compensation are evolutionarily conserved outside of humans, we examined protein expression differences in haploid yeast that had been engineered to harbor single disomic chromosomes^39,40^. We identified one-to-one orthologs between budding yeast and human proteins, and we found a significant correlation between the expression of proteins encoded on aneuploid chromosomes in human cancers and in budding yeast (Figure S3C) (corr = 0.214, P = 4E-09). For instance, the ribosomal subunit RPL38, the splicing factor SF3A1, and the ER membrane complex protein EMC3 exhibited consistent dosage compensation at the protein level when encoded on a chromosome that was gained in cancer cells and when encoded on a chromosome that was gained in yeast (Figure S3D-F). In total, these data demonstrate that the protein expression changes observed on aneuploid chromosomes in a large panel of genomically-diverse cancer cell lines are recapitulated in other aneuploid backgrounds and species, demonstrating that these patterns of scaling and buffering are evolutionarily conserved.

Next we investigated whether mRNA differences upon aneuploidy were similarly conserved. We found that there is a significantly weaker correlation between RNA expression upon chromosome gain in the CCLE dataset, stable aneuploid cell lines, Down syndrome fibroblasts, and aneuploid yeast (Figure S3H-J). The observed correlation coefficients were 1.5 to 3-fold higher at the protein level than at the RNA level (Figure S3K-L). These findings suggest that the conserved patterns of aneuploidy-induced gene expression that we have uncovered are largely controlled at the post-transcriptional level, though some transcriptional similarities are maintained.

### Effects of cellular aneuploidy on global patterns of protein expression

It has previously been reported that aneuploid cells exhibit a set of shared transcriptional changes, independent of the identity of the extra chromosome (e.g., *in trans*^59,60^). To investigate whether aneuploidy is associated with gene expression changes *in trans* across a large panel of cancer cell lines, and to determine where these patterns are maintained at the protein level, we calculated a cellular aneuploidy score based on the total number of genes encoded on aneuploid chromosomes in each cell line, normalized to that line’s basal ploidy. We then performed Pearson correlation analysis to identify genes whose expression was positively or negatively correlated with this cellular aneuploidy score. Positive correlations represent genes that were upregulated in highly-aneuploid cell lines relative to cell lines with lower total aneuploidy, while negative correlations represent genes that were downregulated in highly-aneuploid cell lines (Figure 4A, S4A) (Supplementary Data 3-4).

**Figure 4.**
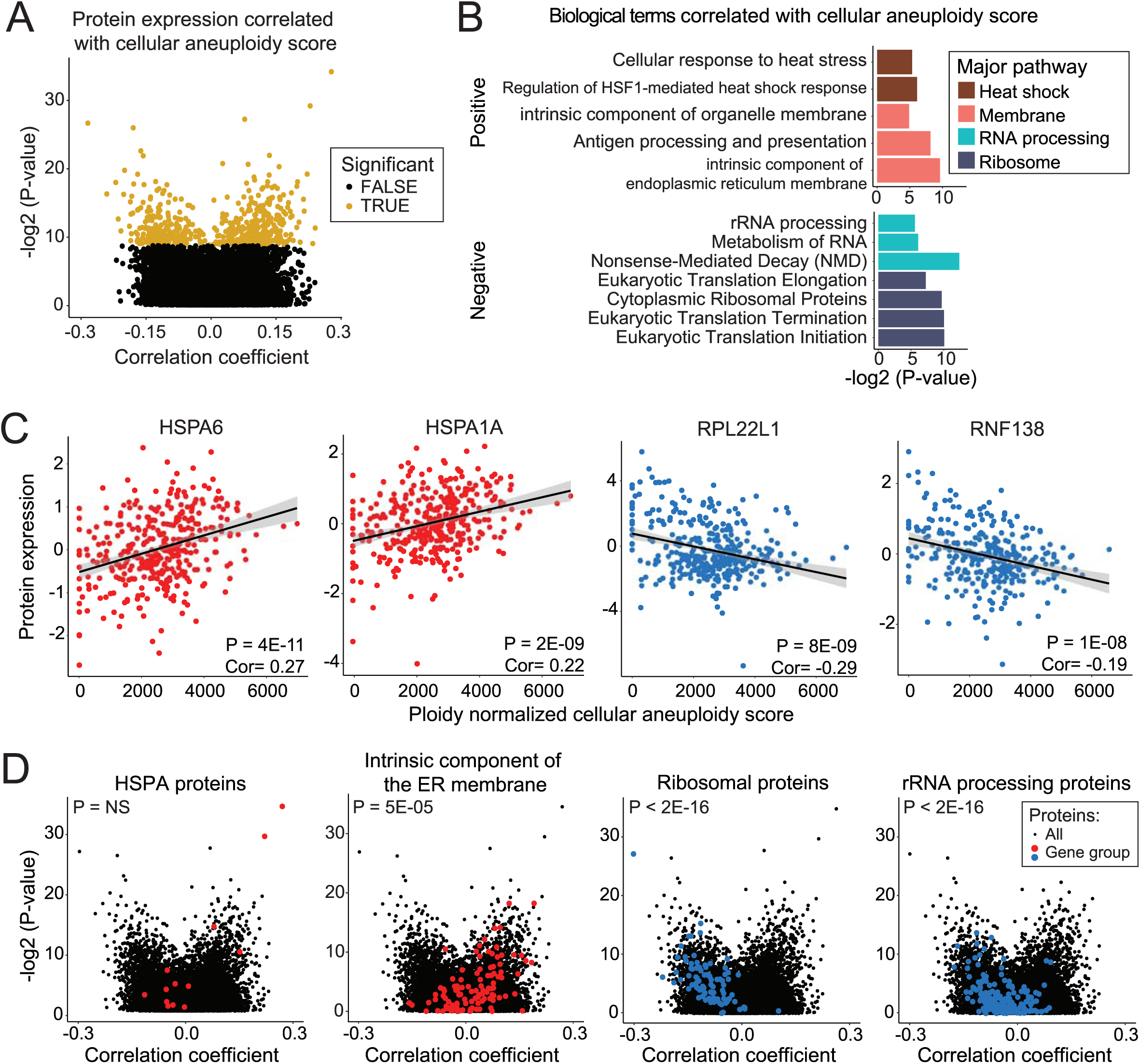
Specific protein expression levels correlate with total cellular aneuploidy. A) A volcano plot displaying the Pearson correlation coefficient between protein expression levels and cellular aneuploidy score vs. the p-value for that comparison. Genes that are significant at a P < 0.05 threshold after Benjamini Hochberg correction with a 5% FDR are labeled in gold. B) A bar graph displaying the GO terms that are enriched among genes positively (top) or negatively (bottom) correlated with total cellular aneuploidy. The complete list of GO terms is included in Supplementary File 3. C) Scatter plots displaying the expression of the two proteins that are most significantly positively correlated with aneuploidy score (left) and the two proteins that are most significantly negatively correlated with aneuploidy score (right). Protein expression is plotted by cellular aneuploidy score. Pearson correlation coefficients and p-values are displayed. Linear regressions and 95% confidence intervals are plotted against the data. D) Volcano plots displaying the correlation coefficient between protein expression and cellular aneuploidy score vs. the corresponding p-value. The background set of all genes are plotted in black, while genes belonging to indicated groups are labeled in red or blue. P-values correspond to two-sided t-tests between correlation coefficients of gene groups and all other genes.

Consistent with prior publications^24,41^, we observed that ribosomal and rRNA processing terms were strongly negatively correlated with cellular aneuploidy scores at both the RNA and protein level (Figure 4B, S4B, Supplementary File 3). Indeed, the two most significantly negatively correlated proteins are the ribosome-associated proteins RPL22L1 and RNF138 (Figure 4C). Another ribosomal gene, RPL3, is the most strongly negatively correlated gene at the RNA level (Figure S4C). The two proteins most positively correlated with aneuploidy scores, HSPA6 and HSPA1A, were both heat shock proteins (Figure 4C). HSPA1A has previously been reported to be upregulated in aneuploid murine fibroblasts^28^. Additionally, genes annotated to “heat assimilation” and “cellular response to heat stress” were enriched among the genes whose expression positively correlated with aneuploidy, which may reflect an aneuploid cell’s increased reliance on protein folding chaperones^36,61^ (Figure 4B, S4B, Supplementary File 3). Finally, membrane-related genes were enriched among genes positively correlated with cellular aneuploidy (Figure 4B). Most notably, intrinsic components of the endoplasmic reticulum (ER) membrane genes were significantly upregulated at the RNA (Figure S4D) and protein level (Figure 4D). Together, these data demonstrate that the expression of certain genes and pathways are associated with the levels of total cellular aneuploidy, independent of whether the genes themselves are encoded on an aneuploid chromosome.

### Both cis and trans effects contribute to dosage compensation in aneuploid cancer cell lines

In the above analysis, we noted that certain gene groups, particularly the ribosome and translation-associated processes, tended to be downregulated in both highly-aneuploid cells and when those genes were encoded on a gained chromosome. We therefore considered the possibility that the “dosage compensation” phenotype that we described above was a consequence of this global response to cellular aneuploidy, rather than a consequence of the altered copy number of these genes *in cis*.

Three analyses indicate that *trans* effects are insufficient to fully account for the protein buffering that we have observed. First, we found that protein complexes and ribosome subunits were upregulated, rather than downregulated, relative to the mean protein when encoded on a lost chromosome (Figure 3B). This change is in the opposite direction of the *trans* effects of aneuploidy, suggesting a chromosome copy number-specific effect. Secondly, we found that the patterns of protein buffering and scaling were maintained in fibroblasts with natural single-chromosome trisomies and congenic cancer cell lines engineered to harbor single extra chromosomes, indicating that cell lines with very low total aneuploidy display dosage compensation of protein complexes (Figure S3). Thirdly, we conducted an additional analysis, in which we isolated and analyzed the quartile of cell lines with the lowest cellular aneuploidy scores and the quartile of cell lines with the highest cellular aneuploidy scores. We then calculated the difference in gene expression upon chromosome arm gain and loss specifically within these quartiles (Supplementary Data 5). We found that ribosome proteins and protein complex subunits were similarly enriched among buffered proteins within both the lowest-aneuploidy quartile and the highest-aneuploidy quartile (Figure S5 A-B, and Supplementary File 4). Thus, while the genome-wide effects of aneuploidy on protein expression can influence the expression landscapes of cancer cell lines, these analyses indicate that dosage effects *in cis* significantly contribute to the buffering phenotype that we have described, independent of total cellular aneuploidy levels.

### Post-translational modifications, protein complex formation, and RNA expression variance contribute to protein buffering upon aneuploidy

We next sought to identify the factors that drive protein dosage compensation. To do this, we calculated the receiver operator characteristic area under the curve (ROC AUC) to estimate a factor’s ability to predict protein buffering. We examined 30 different factors that capture various genetic, biochemical, and biophysical features of each gene or protein, and we assessed their correlation with buffering upon chromosome gain and chromosome loss.

The strongest predictors of buffering upon chromosome gain were the number of ubiquitination sites within a protein, the number of protein-protein interactions that a protein exhibited, and the number of protein complexes a protein is incorporated into (Figure 5A, 5B; AUC = 0.56 - 0.57). The same features were also strongly correlated with buffering upon chromosome loss. Several other post-translational modifications, including protein methylation, phosphorylation, and acetylation, were also correlated with protein buffering upon either chromosome gain or loss. This suggests that protein complex subunits and proteins that are regulated by post-translational modifications tend to exhibit dosage compensation in aneuploid cells.

**Figure 5.**
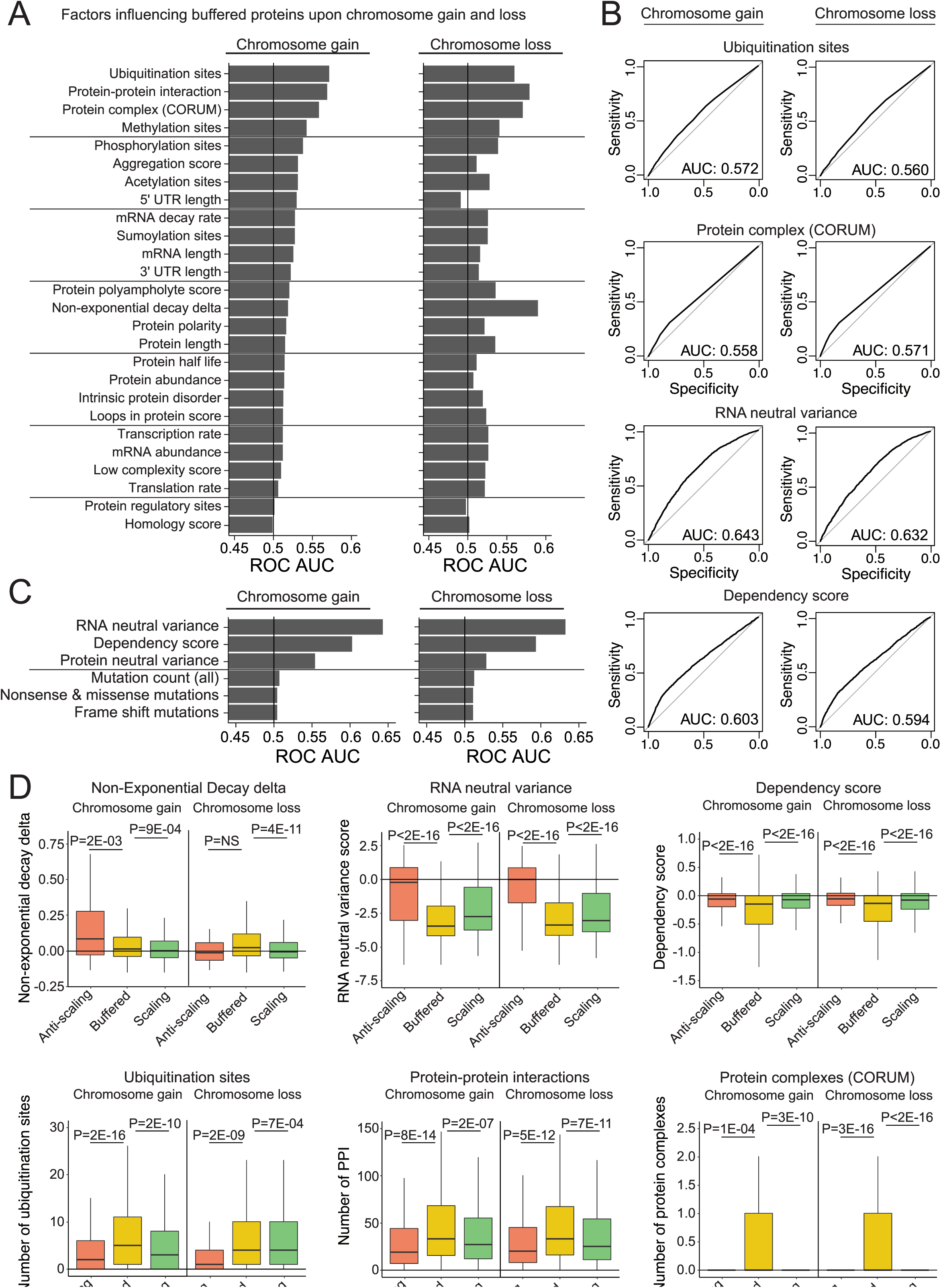
Multiple genetic and biochemical factors predict protein buffering upon aneuploidy. A) Bar graphs displaying ROC area under the curve values for each independent factor upon chromosome gain or loss. Genes classified as “buffered” upon chromosome gain or loss were set as the true positive fraction. B) ROC curves are displayed for certain key factors. Genes classified as “buffered” upon chromosome gain or loss were set as the true positive fraction. C) ROC AUC values for dataset-specific factors upon chromosome gain or loss. Genes classified as “buffered” upon chromosome gain or loss were set as the true positive fraction. D) Boxplots displaying buffering factor scores, per difference category upon chromosome gain or loss. Six buffering factors are displayed: non-exponential decay delta, RNA neutral ploidy variance, dependency score, number of ubiquitination sites, number of protein-protein interactions, and number of protein complexes. P-values are from two-sided t-tests. Boxplots display the 25th, 50th, and 75th percentile of the data, while the whiskers indicate 1.5 interquartile ranges.

Interestingly, several factors associated with protein regulation had minimal ability to predict protein buffering, including protein translation rate, protein half-life, and intrinsic protein disorder. The most notable distinction between factors influencing protein buffering upon chromosome gain and loss was the non-exponential decay (NED) delta, the difference between a protein’s observed degradation kinetics and the expected exponential pattern^42^. NED proteins tend to be initially degraded at a rapid rate, followed by their subsequent stabilization. While the NED delta was a strong predictor of buffering upon chromosome loss (AUC = 0.59), it was less predictive of buffering upon chromosome gain (AUC = 0.52).

We also examined several factors that were specific to this CCLE dataset. We found that buffered proteins exhibited lower expression variation at both the RNA and protein level within cell lines in which that gene was encoded on a non-aneuploid chromosome (Figure 5B, 5C, 5D). This indicates that proteins buffered in aneuploid cells tend to be more tightly regulated, even in euploid conditions. A gene’s dependency score was also found to be a good predictor of protein buffering (Figure 5B, 5C; AUC = 0.59 - 0.60). Finally, we found that the frequency of mutations per gene did not affect their likelihood to be buffered upon aneuploidy, suggesting that nonsense-mediated decay was not a significant factor in dosage compensation.

Next, we examined several of the predictive factors identified above individually. We verified that buffered proteins tended to exhibit lower dependency scores (indicative of essential genes), more ubiquitination sites, and more protein-protein interactions than non-buffered proteins (Figure 5D). In fact, buffered proteins tended to be significantly different from both the scaling and anti-scaling gene groups. For instance, 31% of buffered proteins are members of one or more protein complexes, which was significantly more than we observed among either scaling or anti-scaling proteins. These differences suggest that the anti-scaling gene group is not simply a more extreme instance of protein buffering, and is instead driven by separate factors. Indeed, we observed that anti-scaling genes exhibited significantly higher expression variation when encoded on non-aneuploid chromosomes compared to either buffered or scaling genes. Thus, the unexpected expression changes within this gene group may reflect inherent noise or variability in their regulation.

### Many oncogenes are buffered upon aneuploidy but scale with gene amplification

It is commonly hypothesized that aneuploidy drives tumorigenesis by increasing the expression of OGs and decreasing the expression of TSGs^15,19,20^. To investigate the relationship between aneuploidy and tumorigenesis, we examined the effects of aneuploidy on a set of verified TSGs and OGs^21^. At the individual gene level, some OGs and TSGs significantly scale with chromosome gain and loss (Figure 6A, MAPK1 and SMAD4) while others showed no significant difference in protein expression upon chromosome gain or loss (Figure 6B, MYC and CDK12). We discovered that the average protein expression of OGs and TSGs was not significantly different compared to the background expression of all proteins on aneuploid chromosomes (Figure 6C). Thus, a chromosome gain event increased the expression of the average oncogene by only 12%, and a chromosome loss event decreased the expression of the average tumor suppressor by only 9%. No GO terms were found to be enriched among scaling, buffered, or anti-scaling OGs or TSGs upon chromosome gain or loss.

**Figure 6.**
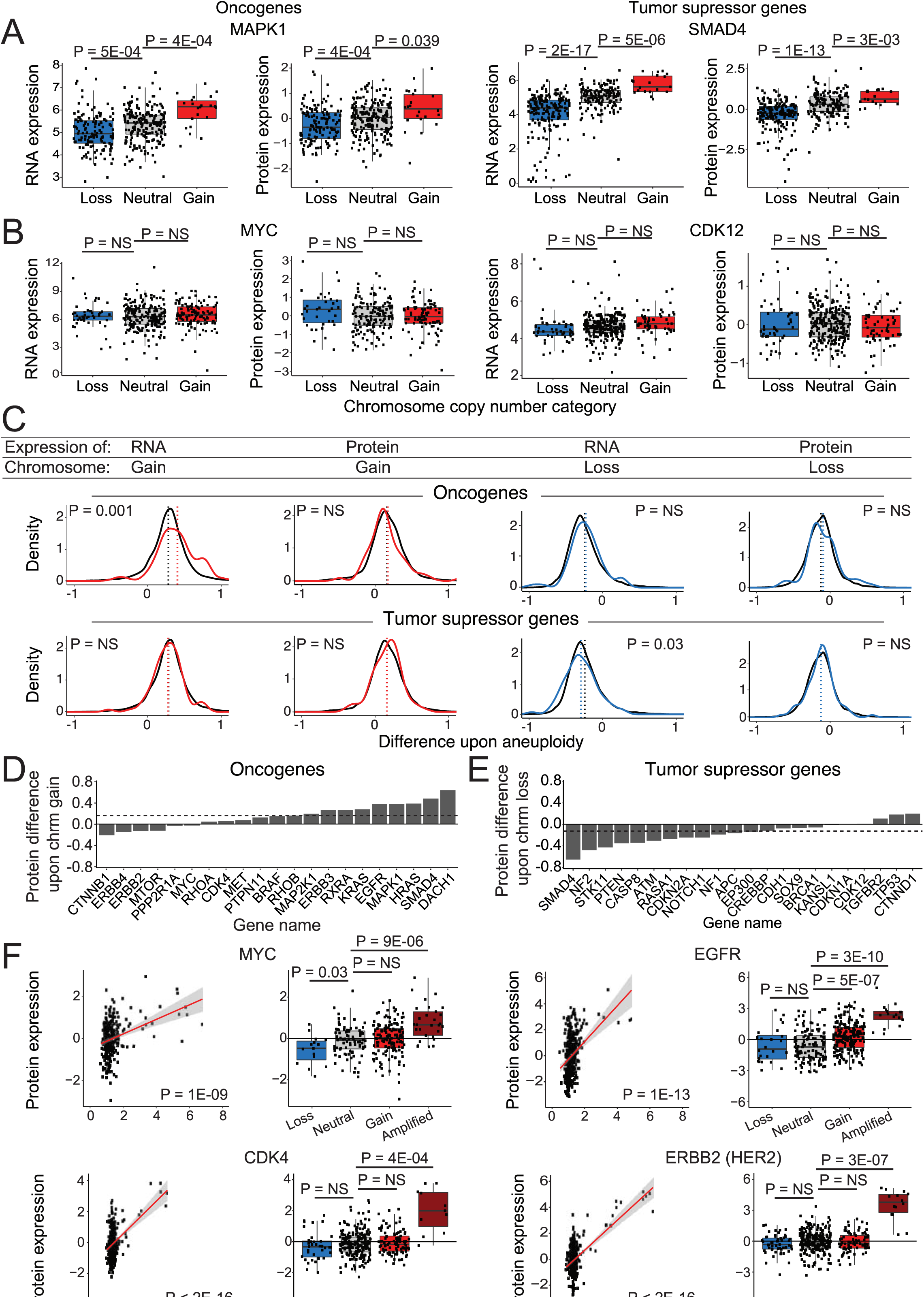
OGs and TSGs can be buffered upon chromosome gain or loss, though some OGs scale with gene copy number amplifications. A-B) Boxplots displaying RNA and protein expression differences upon chromosome gain and loss, for oncogenes (left) and tumor suppressor genes (right). Boxplots display the 25th, 50th, and 75th percentile of the data, while the whiskers indicate 1.5 interquartile ranges. P-values represent two-sided t tests. C) Density curve of OG and TSG expression at the RNA level and protein level upon chromosome gain (red) or chromosome loss (blue). Difference in expression for all other genes is displayed in black. Mean expression differences per condition and gene group are labeled by dotted lines. P-values are from two sided t-tests between the indicated gene group and the background set of all other genes. D-E) Bar graphs displaying the mean protein expression difference for D) OGs upon chromosome gain and E) TSGs upon chromosome loss. Not all oncogenes and TSG are displayed. Dotted lines indicate the mean differences in expression for all genes. Complete list of oncogene and tumor suppressor gene expression differences upon chromosome gain and loss are available in Supplemental Data 2. F) Scatter plots and boxplots displaying protein expression levels for four oncogenes, MYC, EGFR, CDK4, and ERBB2 (HER2), relative to their gene copy number. Linear regressions (red) with 95% confidence intervals (grey) are plotted against the data. P-values in the scatterplots were calculated from Pearson’s correlation coefficient, while p-values in the boxplots were calculated from two-sided t-tests.

Among individual OGs and TSGs, we noted that several potent oncogenes exhibited buffering or anti-scaling upon chromosome gain (MYC, MTOR, ERBB4) (Figure 6D). For instance, the MYC gene is located on chromosome 8q, but we did not detect a significant increase in MYC expression in 110 cell lines with Chr8q gains compared to 211 cell lines that are neutral for Chr8q (Figure 6B). Similarly, the expression of many tumor suppressors, including BRCA1, CDKN1A, and CDH1, did not significantly decrease when the chromosome that they were encoded on was lost (Figure 6E).

If oncogenes are subjected to dosage compensation in aneuploid cells, how could increases in oncogene expression levels arise to drive tumorigenesis? We observed that several oncogenes that were subject to dosage compensation upon single-chromosome gains were nonetheless sensitive to gene-level focal amplifications. For example, 11% of cancer cell lines exhibited focal amplifications of the MYC locus, and this was associated with a significant 86% (0.895 log2 fold change) increase in MYC protein expression (Figure 6F). Similar results were obtained for other potent OGs, including EGFR, CDK4, and ERBB2 (HER2) (Figure 6F). Thus, oncogene dosage compensation is imperfect. While genes like MYC can be resistant to expression changes resulting from arm-length aneuploidies, focal amplifications overcome these compensatory mechanisms and cause an increase in driver oncogene expression.

## Discussion

Here, we conduct a genome-wide analysis of protein expression changes in aneuploid cancer cell lines. We show that, while protein expression tends to increase upon chromosome gain, and protein expression tends to decrease upon chromosome loss, a majority of proteins are subjected to dosage compensation. These patterns of scaling and buffering upon aneuploidy are conserved between aneuploid yeast cells, primary human fibroblasts, and immortalized cancer cell lines. Dosage compensation in aneuploid cancers may counteract alterations in the expression of oncogenes and tumor suppressors, and therefore a comprehensive understanding of this phenomenon will shed light on how genomic alterations drive tumorigenesis.

Consistent with previous results^41,42,62,63^, we found that protein complex subunits exhibited significant buffering when they were encoded upon a gained chromosome arm. This trend is exemplified by ribosomal complex proteins^24,26^, which showed highly-significant dosage compensation. Interestingly, we discovered that protein complex subunits were also significantly buffered upon chromosome arm loss. Protein complex RNA levels tend to scale with chromosome copy number, indicating that buffering occurs primarily at the post-transcriptional level. We hypothesize that the stabilization and reduced degradation that proteins experience when they are incorporated into complexes may contribute to expression buffering upon chromosome loss, while the rapid degradation^42^ of unincorporated subunits may contribute to protein buffering upon chromosome gain. In support of this hypothesis, non-exponential decay kinetics^42^ are more predictive for buffered proteins upon chromosome loss than gain. Thus, cells may have unique pathways that are able to maintain appropriate complex stoichiometries in the event of both subunit overproduction and subunit underproduction.

Our study has several limitations. Notably, our analysis was performed across genetically-diverse human cell lines derived from a variety of cancer lineages. These cancers exhibit varying degrees of aneuploidy, which are also capable of affecting gene expression *in trans* through the induction of aneuploidy-associated stresses^59,60^. Additionally, cancers may display a number of sub-chromosomal alterations, including focal amplifications and deletions, that were not isolated in this analysis^64,65^. Nonetheless, we observed striking expression similarities between our genome-wide pan-cancer analysis and congenic cell lines engineered to harbor single extra chromosomes, primary fibroblasts naturally trisomic for chromosome 21, and even yeast strains harboring additional yeast chromosomes. These findings indicate that, in spite of the genetic diversity and potential presence of subchromosomal alterations, the patterns of buffering and scaling that we describe reflect protein dosage changes that are shared in other aneuploid conditions. Moreover, the evolutionary conservation of these buffering patterns may reflect highly-conserved methods for controlling the stoichiometry of protein complex subunits. Interestingly, the conservation of protein expression patterns between aneuploid cancers and Down syndrome fibroblasts suggests that the patterns we have described could also contribute to aneuploidy-associated developmental syndromes^66–68^.

Several distinct genetic and biochemical factors were capable of predicting protein buffering. In addition to protein-protein interactions, these factors included a number of post-translational modifications, aggregation propensity, and gene essentiality. Unexpectedly, we also discovered that 10% of proteins encoded on an aneuploid chromosome exhibited an opposite directional change relative to the chromosome copy number alteration. Our analysis highlights that buffered proteins and these “anti-scaling” proteins are differentially regulated by a variety of biological factors. For example, anti-scaling genes are less likely to be part of a protein complex than buffered genes and exhibit fewer total protein-protein interactions. Additionally, anti-scaling genes exhibit a very high neutral ploidy expression variance, while buffered genes exhibit low neutral ploidy expression variance. The anti-scaling gene group was highly enriched for extracellular components, and more research will be needed to fully elucidate the regulation of these genes and their effects on the physiology of aneuploid cancers.

It is commonly hypothesized that one of the main drivers of the evolution of aneuploid cancers is the copy number of oncogenes and tumor suppressors^19^. While many oncogenes exhibit increased expression upon chromosome gain, and many tumor suppressors exhibit decreased expression upon chromosome loss, we discovered that this is not true for all oncogenes and tumor suppressors. Indeed, many important oncogenes, including HER2, MTOR, and MYC, do not increase in expression with the gain of the corresponding chromosome arm, and the overall effect of chromosome gains across all oncogenes was moderate. Notably, this analysis was performed in immortalized cancer cell lines, and therefore may not reflect expression patterns immediately after chromosome missegregation and/or early in tumor development. Further research conducted on *in vivo* specimens and following induced aneuploidy events may shed light on the immediate effects of chromosome gains and losses. However, we note that cancer cell lines harboring focal amplifications of key oncogenes including MYC and HER2 exhibited highly-significant increases in the protein expression of these genes. Thus, the buffering of oncogenes that we have observed is imperfect, which may promote the evolution of high-copy amplifications to increase oncogene expression and escape compensatory downregulation.

Oncogene and tumor suppressor gene dosage compensation complicates, but does not contradict, the aneuploidy/driver gene dosage hypothesis. Developing a comprehensive picture of oncogene and tumor suppressor buffering will improve our understanding of cancer driver genes and their role in shaping aneuploid karyotypes in cancer. As aneuploidy has been recognized to be a highly-significant prognostic factor across cancer types^3,13–18^, and as genomic and exomic sequencing costs have rapidly fallen^69^, we expect that high-resolution cancer karyotyping will become increasingly routine in a clinical setting. Knowing which proteins are sensitive to chromosome gains and losses may help identify the specific drivers in a patient’s tumor, leading to better personalized therapies and improved patient outcomes.

## Methods

### Data sources

The effect of aneuploidy on gene expression was analyzed using DNA, RNA and protein data generated through the Cancer Cell Line Encyclopedia and available in processed form from the DepMap project^49–54^.

*Protein expression data* was retrieved from the DepMap “Proteomics” dataset (downloaded August 3^rd^, 2020). The protein data had been normalized, log2 transformed, and mean expression was set at 0^49,50^.

*RNA expression data* was retrieved from the DepMap “Expression” (Public 20Q4) dataset^51,53^. The RNA data was log2 transformed with a pseudocount of 1.

“*Cell Line Sample Info”* was downloaded from DepMap (May 21^st^ 2020) and used to link various data types^51^.

*Relative gene copy number data* was retrieved from DepMap “CCLE gene cn” (Public 21Q1) dataset on April 29, 2021^52^. The data was log2 transformed with a pseudocount of 1.

*Chromosome arm copy number data* were retrieved from Cohen-Sharir, Y et al. 2021^55^.

*Stable aneuploidy cell line data* were retrieved from Stingele et al. 2012^24^. Protein and RNA difference upon aneuploidy was calculated by taking the log2 fold change between aneuploidy clones and their near-euploid cell line controls. For protein data, the mean difference was taken for all four cell lines with a gain for chromosome 5: RPE-1 trisomy 12 and 5, HCT-116 tetraploidy 5, HCT-116 H2B-GFP tetraploidy 5 and HCT-116 H2B-GFP trisomy 5. For RNA data, mean RNA difference was found from RPE-1 trisomy 12 and 5 and HCT-116 tetraploidy 5, the only two chromosome 5 gain transcriptomes available. Mean gene expression difference was plotted against DepMap protein difference data.

*Down syndrome aneuploidy data* was retrieved from Liu et al.^57^ and Letourneau et al.^58^. Protein expression difference was calculated by taking the log2 fold change in quantile-normalized protein expression from 11 Down syndrome fibroblast lines and matched controls. The difference in RNA expression was reported as the log2 fold change in RNA expression between a pair of monozygotic twins discordant for chromosome 21.

*Yeast aneuploidy gene expression difference* was retrieved from Dephoure et al. (2014)^39^. Protein and RNA expression differences were reported as log2 ratios for genes located on duplicated chromosomes.

*Oncogene and tumor suppressor gene lists* were acquired from Bailey et al. (2018)^21^ based on a comprehensive analysis of TCGA data.

*Potential buffering factors* explored in Figure 5 were retrieved from multiple sources. Several potential buffering factors were retrieved from MobiBD (Version: 4.0 - Release: 2020_09^70^). MobiBD data includes: *Protein intrinsic disorder* (prediction-disorder-mobidb_lite), *low complexity score* (prediction-low_complexity-merge), *homology score* (homology-domain -merge), *loops in protein score* (prediction-lip -anchor), *protein polyampholyte score* (prediction-polyampholyte-mobidb_lite_sub) and *protein polarity* (prediction-polar-mobidb_lite_sub).

*Translation rate, transcription rate, protein length, mRNA length, mRNA abundance and protein abundance* data was retrieved from Hausser et al. 2019^71^. *Protein half life* data was from Methieson et al. 2018^72^, the mean protein half life was used. *mRNA decay rates* were retrieved from Yang et al. 2003^73^.

*UTR data (5’ and 3’)* was retrieved from the UCSC human genome browser (hg38 GRCh38). The difference between transcription start sites and coding region start sites was found for 5’ UTR, and the difference between coding sequence end and transcription sequence end was found for 3’ UTR regions. Only manually curated (“NM”) mRNA genes were analyzed.

Datasets for protein *acetylation, methylation, phosphorylation, ubiquitination, sumoylation and gene regulatory sites* were downloaded from phosphosite plus (Hornbeck et al. 2015^74^) (last updated on April 19, 2021). The number of regulatory and/or modification sites were found per human gene. Genes not found in the dataset were set to zero modification sites.

The *neutral variance for RNA and protein expression* was calculated from DepMap protein and RNA expression data. Cell lines without a gain or loss (neutral ploidy) of the chromosome arm a gene was located on were used to calculate the neutral variance for that gene. The coefficient of variance was taken from gene expression levels and log2 transformed.

The *mutation counts (all)* was calculated per gene using the DepMap CCLE mutations public dataset (21Q2, Broad 2021^51^). The mutation data was filtered for the cell lines in our expression difference dataset, and the number of all mutations was counted per gene. For the *mutation counts (nonsense and missense)*, as well as the *mutation counts (frame shifts)* only counted mutations of the indicated variants.

Non-exponential decay delta scores were taken from McShane et al. 2016^42^.

The *aggregation score* per protein was extracted from Ciryam et al. 2013^75^.

*Protein complex (CORUM) score* was extracted from the comprehensive resource of mammalian protein complexes (CORUM) “complete complex” dataset, version 3.0 (last updated 2018)^76^. All human protein complexes were extracted, and the number of times a protein appeared in the dataset was calculated. Proteins not found in protein complexes were given a score of zero.

The number of *protein-protein interactions* per protein was extracted from the Human Integrated Protein-Protein Interaction rEference (HIPPIE), version 2.2 (last updated February 2019)^77^. Interactions with confidence values >0.6 were used, and the sum of interactions per protein was calculated. Proteins not found to interact with other proteins were given a score of zero.

*Dependency scores* were extracted from DepMap Achilles gene dependency (21Q2^52^) dataset. We took the mean gene dependency from across all reported cell lines.

### Dataset filtering

We generated a *dataset measuring gene expression difference upon chromosome gain or loss*. We only used cell lines with proteomics data, RNA expression data and chromosome arm copy number data (367 cell lines). We only included a gene in our difference analysis if we had RNA and protein expression data for that gene from at least 10 cell lines in which the chromosome arm in which that gene was encoded on was gained, 10 cell lines in which the chromosome arm was lost, and 10 cell lines in which the chromosome arm had a neutral ploidy.

We generated two datasets measuring *gene expression differences upon chromosome gain and loss within cells with either low or high levels of cellular aneuploidy*. As above, we initially included cell lines with proteomics data, RNA expression data, and chromosome arm copy number data. Next, we isolated the quartile of the cell lines with the lowest cellular aneuploidy score (from 0 to 1773), and the quartile of cell lines with the highest cellular aneuploidy score (from 3309 to 6985). We only included a gene in our difference analysis if we had RNA and protein expression data for that gene from at least 3 cell lines per chromosome arm category, as described above (Supplementary Data 5).

For the *protein expression correlation with aneuploidy score dataset* used for Figure 4, all cell lines with protein expression and chromosome arm copy number data were used for protein aneuploidy score analysis. Similarly, for the *RNA expression correlation with aneuploidy score* dataset used in Supplementary Figure 4, all cell lines with RNA expression measurements and chromosome arm copy number data were used.

### Analyzing the effects of aneuploidy on gene expression

*Gene expression difference upon aneuploidy* was calculated for each gene, for both RNA and protein data. For each gene, the chromosome arm the gene was located on was identified. Next, cell lines in the CCLE filtered dataset (see above) were grouped as either having a chromosome arm gain, chromosome arm loss, or having a neutral ploidy for that chromosome arm. Then, the mean RNA and protein expression for the gene of interest was taken per DNA aneuploidy category (gain, loss, or neutral). The mean gene expression in the neutral category was subtracted from the mean gene expression in the chromosome gain category to obtain the difference in expression upon chromosome gain. Similarly, mean gene expression in the neutral category was subtracted from the mean protein expression in the chromosome loss category to get the difference in expression upon chromosome loss. We found the difference in expression upon chromosome gain and loss for both protein expression data and RNA expression data. As protein and RNA expression data have already been log2 transformed, the difference (log2(A) - log2(B)) is equivalent to log2 fold change (log2(A/B)). P-values per gene were calculated by using a two-sided t-test between gene expression in cell lines neutral for the corresponding chromosome arm and cell lines with either a chromosome gain or loss (Supplementary Data 2, 5).

*Cell line specific difference* was calculated for RNA and protein data in Figure 1 and Figure S1. The mean RNA and protein expression per chromosome arm was taken from cell lines neutral for chromosome arm gains or losses. This “neutral” gene expression mean was subtracted from the mean gene expression per chromosome arm in the cell line of interest.

*Gene expression categories* were based on difference upon chromosome gain or loss in RNA or protein expression data. For chromosome gain events, genes were classified as “anti-scaling” if their mean expression difference was less than -0.1 relative to cell lines that were neutral for that chromosome arm. Genes were classified as “buffered” if their mean expression ranged from -0.1 to 0.25 relative to cell lines that were neutral for that chromosome arm. Genes were classified as “scaling” if their mean expression was greater than 0.25 relative to cell lines that were neutral for that chromosome arm. For chromosome loss events, the values were reversed (> 0.1 for anti-scaling, between 0.1 and -0.25 for buffered and < -.25 for scaling).

*Aneuploidy score*s were calculated by summing the number of genes on all aneuploid chromosome arms in a cell line and then dividing that total by the cell’s basal ploidy. In this way, the gain or loss of a chromosome arm contributes equally to the aneuploidy score.

*Aneuploidy score correlation* was calculated per gene for both protein expression and RNA expression data. Pearson’s correlations between protein or RNA expression and a cell’s aneuploidy score were measured for every gene. Associated p-values are Pearson correlation significance measurements (Supplementary Data 3-4).

### Assessing the conservation of aneuploidy-associated dosage compensation

Gene expression datasets from stable human aneuploidy cell lines, Down syndrome fibroblast lines, and aneuploid yeast strains were acquired and processed as described above. The yeast orthologs of human genes were identified using g:Profiler^56^. For this analysis, only one-to-one orthologs were considered. That is, we only included a yeast gene if it had a single human ortholog and if that human gene only had a single yeast ortholog. Dosage compensation across these various aneuploidy conditions was compared using Pearson correlation coefficients.

### GO term enrichment analysis

*Biological term enrichments* were identified using g:Profiler^56^. Significance values were calculated against a custom background set of genes or proteins that were included in our processed and filtered dataset.

### Identifying genomic features that correlate with protein buffering

*ROC plots and area under the curve analysis were* performed using R package “pROC” version 1.17.0.1^78^ and the various genetic, biochemical, and biophysical datasets listed above. A gene’s classification as “buffering” was used as the true positive “sensitivity” fraction.

### Oncogene and tumor suppressor buffering analysis

*Relative gene copy number categories* were classified as “loss” with a score <0.9, “neutral” at 0.9-1.1, “gain” at 1.1-1.75 and “amplified” at 1.75+. Relative gene copy numbers are log2-transformed data, with a pseudo-count of 1.

### Code availability

The original code used for data analysis has been deposited on GitHub at https://github.com/kschukken/CCLE_protein_dosage_compensation.git. All code was written in R.

## Supporting information

Supplemental File 1. GO enrichment among protein categories

Supplemental File 2. GO enrichment among RNA categories

Supplemental File 3. GO enrichment per total aneuploidy correlation

Supplemental File 4. GO enrichment within low-aneuploidy and high-aneuploidy cell lines

Supplemental Data 1. Cell line information

Supplemental Data 2. Expression differences upon chromosome gain and chromosome loss

Supplemental Data 3. Correlation between protein expression and total cellular aneuploidy

Supplemental Data 4. Correlation between RNA expression and total cellular aneuploidy

Supplemental Data 5. Expression differences upon chromosome gain and chromosome loss in low-aneuploidy and high-aneuploidy cell lines

Supplemental Data 6. Lists of genes per gene groups

## Acknowledgments

Research in the Sheltzer Lab is supported by an NIH Early Independence award (1DP5OD021385), NIH grant R01CA237652, Department of Defense grant W81XWH-20-1-068, a Damon Runyon-Rachleff Innovation award, an American Cancer Society Research Scholar Grant, and a grant from the New York Community Trust.

## Author Contributions

K.M.S. and J.M.S. conceived, designed, and performed the analysis described in this work.

K.M.S. and J.M.S. wrote the manuscript and prepared the figures.

## Legends

**Supplementary Figure 1.**
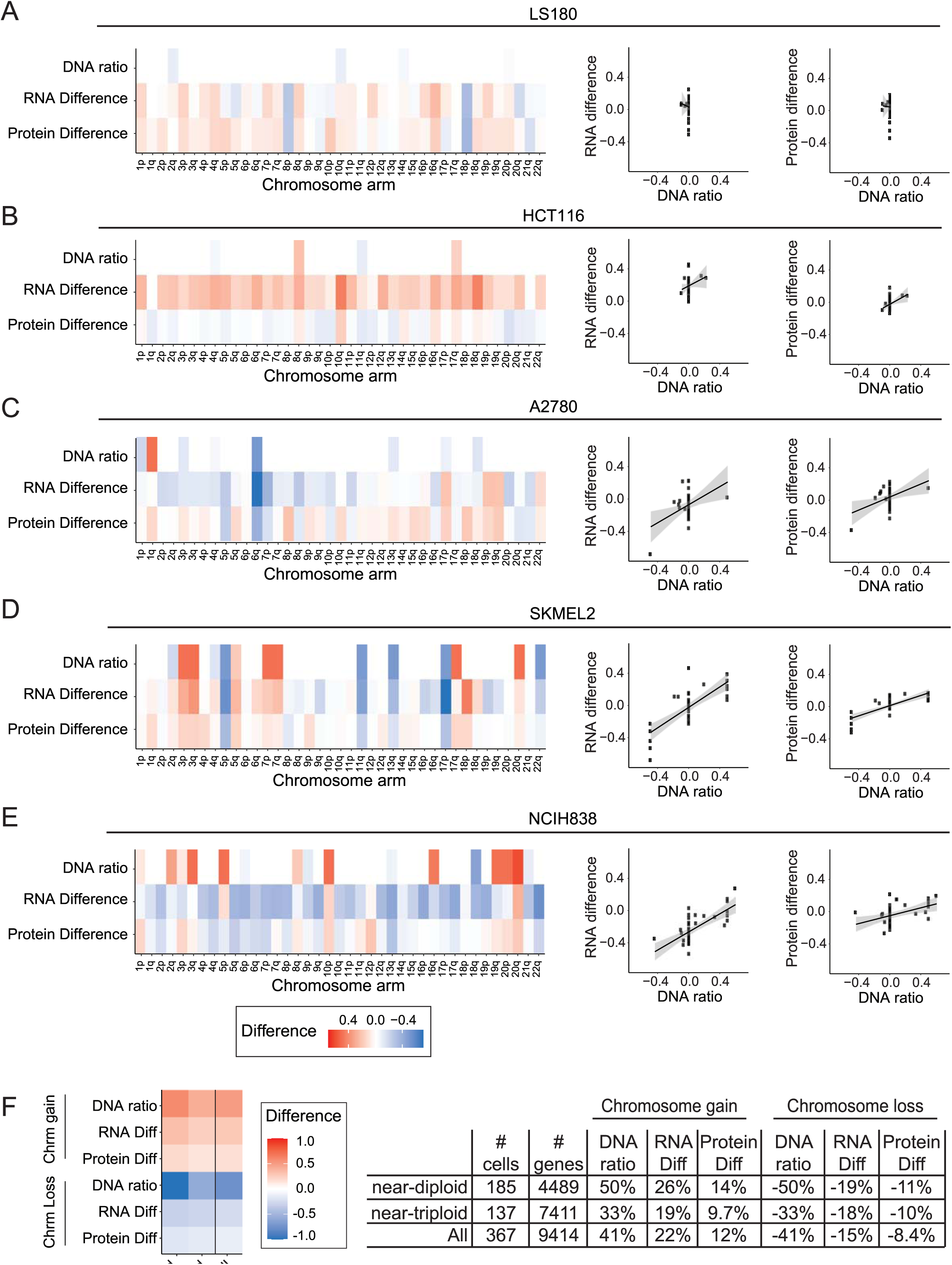
RNA and protein expression differences within individual cancer cell lines and expression differences split based on cell line ploidy. A-E) Heatmap of the DNA ratio, RNA expression difference, and protein expression difference, per chromosome arm, in the indicated cell lines relative to cells neutral for that chromosome arm (left). Scatterplots showing the relationship between the DNA ratio and the RNA or protein expression differences (right). A linear regression with 95% confidence intervals is plotted against the data. F) Heatmap of the DNA ratio, and the difference in RNA and protein expression per chromosome arm, according to the cell line’s basal ploidy (left). Table displaying the quantification of mean difference in RNA or protein expression upon chromosome gain or loss, per ploidy (right).

**Supplementary Figure 2.**
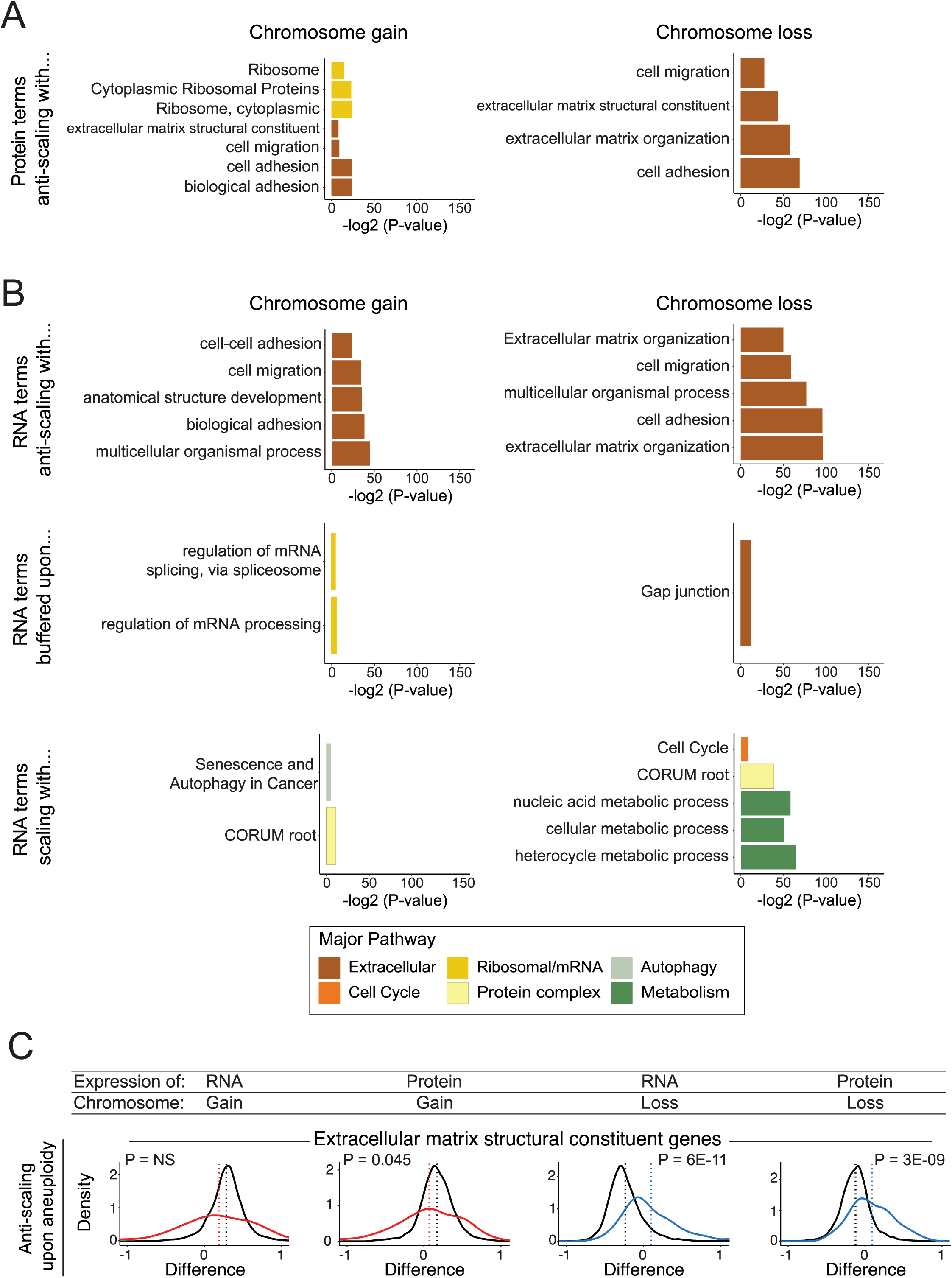
GO term enrichment analysis of anti-scaling proteins and RNA expression categories. A) Bar graphs displaying the GO terms enriched among proteins that exhibit anti-scaling upon chromosome gain or loss. The complete set of GO terms is included in Supplementary File 1. B) Bar graphs displaying the GO terms enriched among RNAs that exhibit the indicated expression changes upon chromosome gain or loss. The complete set of GO terms is included in Supplementary File 2. C) Density graphs of the difference in RNA or protein expression per gene group upon chromosome gain (red) or loss (blue) for extracellular matrix structural constituent genes. Difference in expression for all other genes shown in black. Mean difference per gene group (red or blue), and all other genes (black), is indicated by the dotted lines. P-values are from two-sided t-tests between the indicated gene group and the background set of all other genes.

**Supplementary Figure 3.**
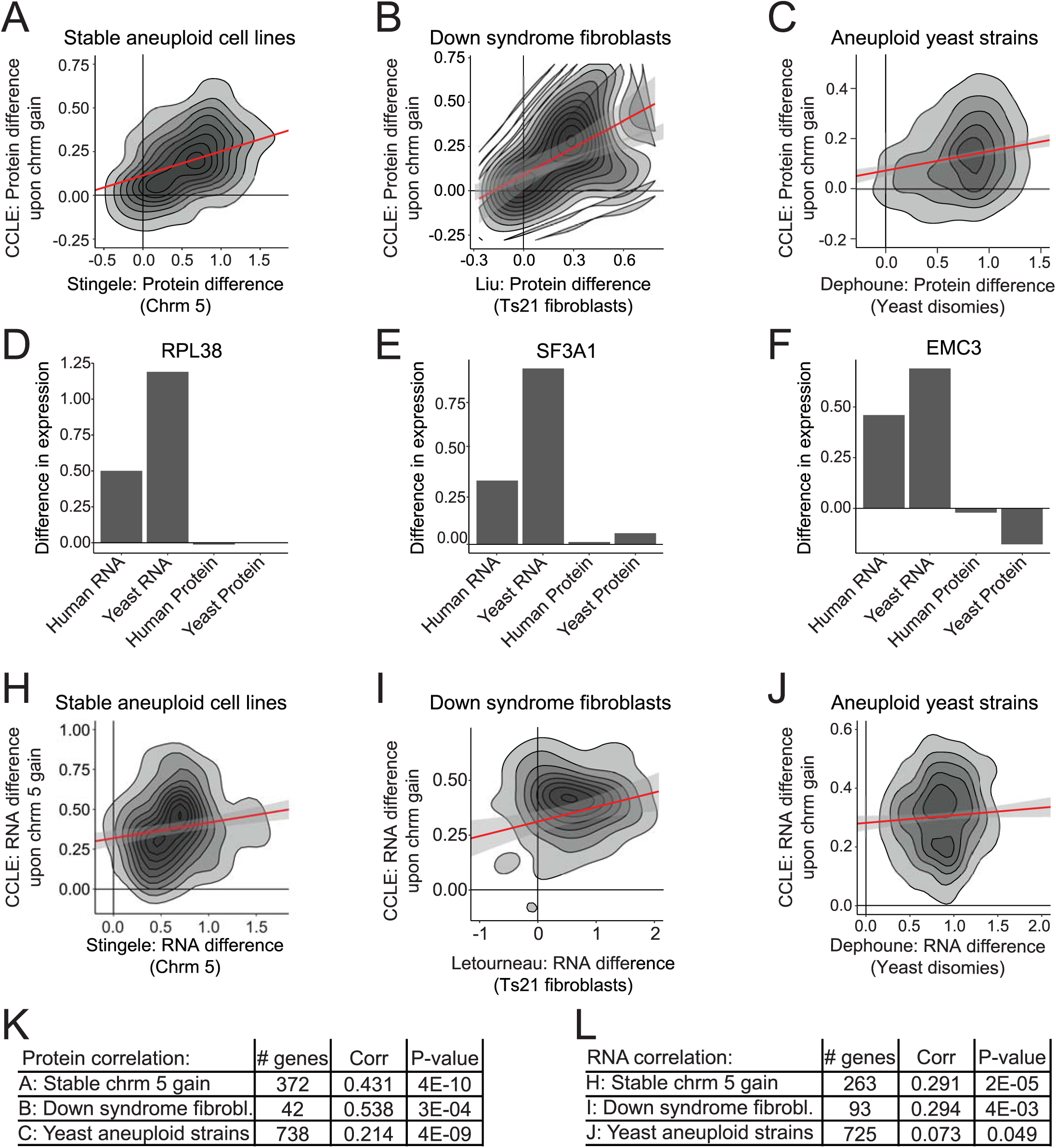
Patterns of protein buffering and scaling are evolutionarily conserved. A-C) Two-dimensional density plots displaying the difference in protein expression upon chromosome gain, for genes encoded on that respective chromosome, in the CCLE dataset versus A) stable aneuploid cell lines with a gain for chromosome 5, B) Down syndrome fibroblasts normalized to matched euploid controls, and C) haploid yeast with single chromosome disomies. Linear correlations (red) with 95% confidence interval (grey) are plotted against the data. D-F) Bar graphs displaying the difference in protein or RNA expression in the CCLE dataset and aneuploid yeast, upon gain of the corresponding chromosome for three genes: D) RPL38, E) SFA1, and F) EMC3. H-J) Two-dimensional density plots displaying the difference in RNA expression upon chromosome gain, for genes encoded on that respective chromosome, in the CCLE dataset versus H) stable aneuploid cell lines with a gain for chromosome 5, I) Down syndrome fibroblasts normalized relative to matched euploid controls, and J) haploid monosomic yeast with single chromosome disomies. Two dimensional density plots with linear correlations (red) with 95% confidence interval (grey) are plotted against the data. K-L) Tables indicating the number of genes analyzed per condition, Pearson’s correlation coefficients, and corresponding p-values for protein expression differences examined in A-C and RNA expression differences examined H-I.

**Supplementary Figure 4.**
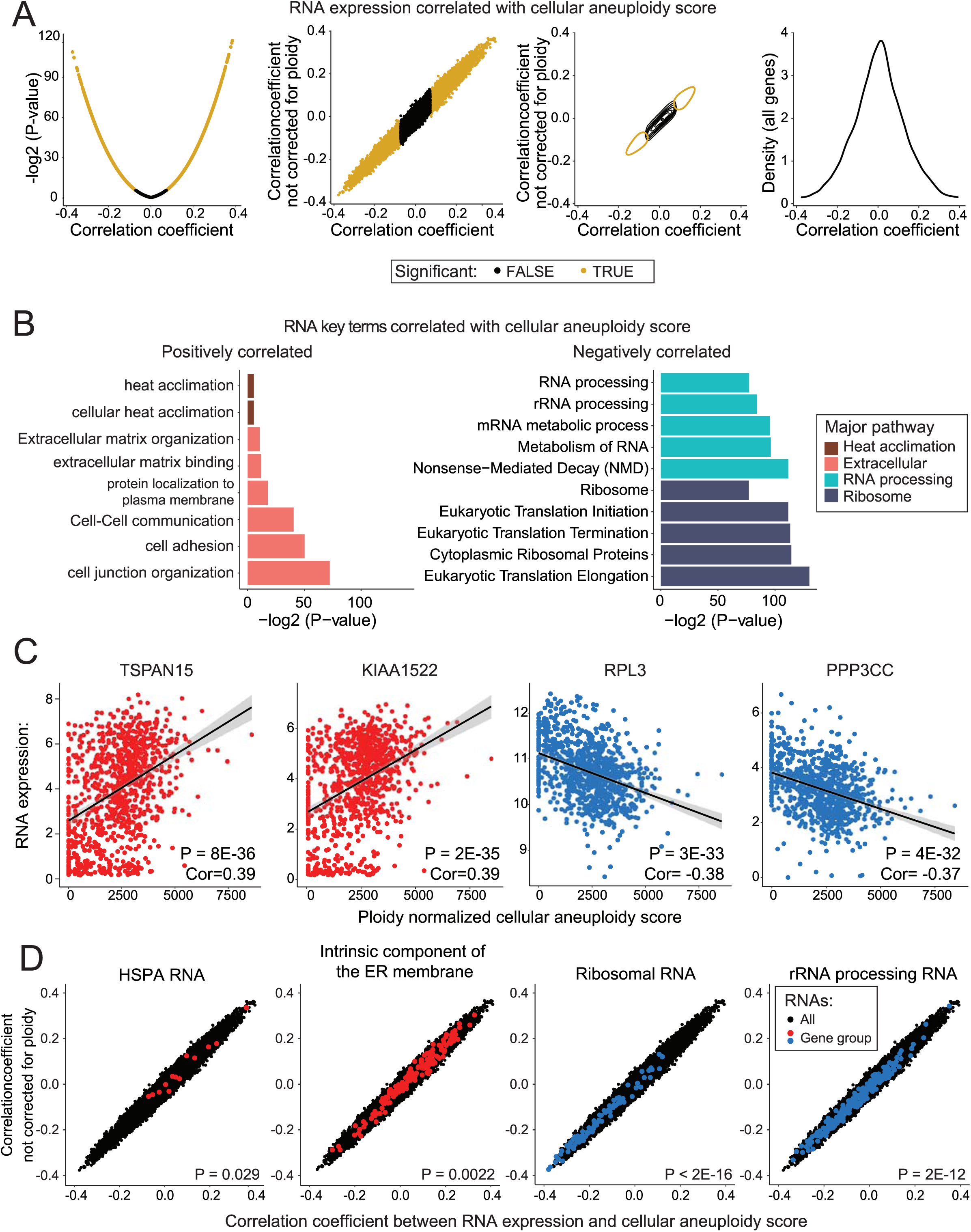
Specific RNA expression levels correlate with total cellular aneuploidy scores. A) Pearson’s correlation coefficients between RNA expression levels and cellular aneuploidy score were calculated. To better view the distribution of this data, the data was plotted in four ways. The negative log2 of the uncorrected p-value was set relative to the correlation coefficient (left). Correlation coefficient between RNA expression and either cellular aneuploidy score (x axis), or the cellular aneuploidy score not corrected for cellular ploidy (y axis) was plotted in two ways (center two graphs). Finally, a one dimensional density plot of the correlation coefficient between RNA expression and cellular aneuploidy score (right). Genes that are significant at a P < 0.05 threshold after Benjamini Hochberg correction with a 5% FDR are labeled in gold. B) Bar graphs displaying the GO terms enriched among RNAs that are positively or negatively correlated with cellular aneuploidy score. Terms are color-coded by their overarching pathway. C) The two RNAs that are most significantly positively correlated (left, red, TSPAN15 and KIAA1522), and negatively correlated (right, blue, RPL3 and PPP3CC) with aneuploidy scores are displayed. RNA expression is plotted according to the aneuploidy score per cell line. D) Scatter plots displaying the correlation coefficients between gene expression and cellular aneuploidy score and the cellular aneuploidy score not corrected for ploidy are displayed for all genes (black), as well as the indicated gene groups (red or blue). HSPA genes and “intrinsic component of the ER membrane” genes (red) are significantly more positively correlated with cellular aneuploidy scores than other genes. Ribosomal RNAs and rRNA processing genes (blue) are significantly more negatively correlated with aneuploidy scores than other genes. P-values from two-sided t-tests of RNA expression in gene groups vs all other genes are displayed.

**Supplementary Figure 5.**
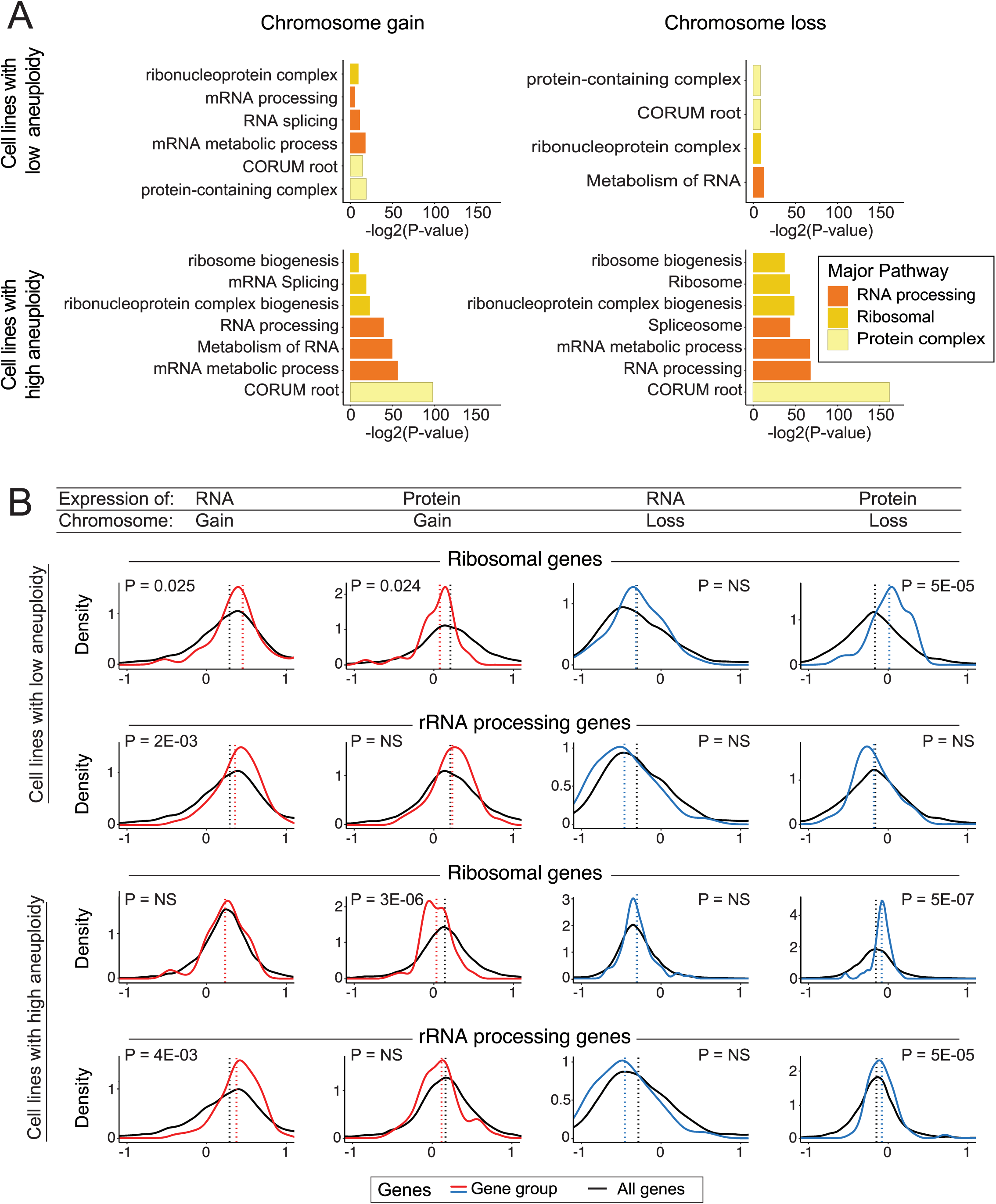
The ribosome and protein complex subunits are buffered in cell lines with low total aneuploidy. A) A bar graph displaying the GO terms that are enriched among buffered proteins within the quartile of cell lines with the lowest total aneuploidy scores (top) and the highest total aneuploidy scores (bottom). The complete list of GO terms is included in Supplementary File 4. B) Density graphs displaying the difference in RNA or protein expression per gene group upon chromosome gain (red) or loss (blue). The difference in expression for all other genes is shown in black. The mean difference per gene group is indicated by the dotted lines. P-values represent two-sided t-tests between the indicated gene groups and the background set of all other genes.

